# Metagenome-derived virus-microbe ratios across ecosystems

**DOI:** 10.1101/2021.02.17.431581

**Authors:** Purificación López-García, Ana Gutiérrez-Preciado, Mart Krupovic, Maria Ciobanu, Philippe Deschamps, Ludwig Jardillier, Mario López-Pérez, Francisco Rodríguez-Valera, David Moreira

**Author notes:** These authors contributed equally to this work (Purificación López-García, Ana Gutiérrez-Preciado).

## Abstract

It is generally assumed that viruses outnumber cells on Earth by at least tenfold. Virus-to-microbe ratios (VMR) are largely based on counts of fluorescently labelled virus-like particles. However, these exclude intracellular viruses and potentially include false positives (DNA-containing vesicles, gene-transfer agents, unspecifically stained inert particles). Here, we develop a metagenome-based VMR estimate (mVRM) that accounts for DNA viruses across all stages of their replication cycles (virion, intracellular lytic and lysogenic) by using normalised RPKM (reads per kilobase of gene sequence per million of mapped metagenome reads) counts of the major capsid protein (MCP) genes and cellular universal single-copy genes (USCGs) as proxies for virus and cell counts, respectively. After benchmarking this strategy using mock metagenomes with increasing VMR, we inferred mVMR across different biomes. To properly estimate mVMR in aquatic ecosystems, we generated metagenomes from co-occurring cellular and viral fractions (>50 kDa-200 µm size-range) in freshwater, seawater and solar saltern ponds (10 metagenomes, 2 control metaviromes). Viruses outnumbered cells in freshwater by ∼13 fold and in plankton from marine and saline waters by ∼2-4 fold. However, across an additional set of 121 diverse non-aquatic metagenomes including microbial mats, microbialites, soils, freshwater and marine sediments and metazoan-associated microbiomes, viruses, on average, outnumbered cells by barely two-fold. Although viruses likely are the most diverse biological entities on Earth, their global numbers might be closer to those of cells than previously estimated.

## Introduction

Viruses are strict molecular parasites that play crucial roles in ecosystems. They control host population sizes and contribute to biogeochemical nutrient cycling [1, 2], helping to maintain biodiversity through *“kill-the-winner”* strategies [3, 4]. They are also important players in evolution, exerting selective pressures on cell populations and triggering arms-race processes, fostering the evolution of genes and genomes and serving as vehicles for horizontal gene transfer [5]. Viruses are also enormously diverse, with new lineages being widely uncovered by metagenomic and metatranscriptomic approaches [6, 7]. It seems therefore natural to think that viruses are also very abundant, but their actual global number and how it compares to that of their hosts remains unclear.

Viruses are generally thought to be at least one order of magnitude more abundant than microbial cells on Earth [2, 8–10]. These abundance estimates are largely based on epifluorescence microscopy or flow cytometry (FCM) counts of virus-like particles (VLPs) and cells stained with nucleic acid dyes. These methods were originally applied to marine plankton, where measured virus-to-microbe ratios (VMR) frequently revolve, despite substantial variation, around 10 (Ref. [1, 9, 11]). However, epifluorescence-based VLP counts exclude temperate viruses and are blind to intracellular lytic viruses during the eclipse phase of infection. Conversely, these methods can suffer from diverse false positives, including cell-DNA-containing membrane vesicles [12–15] or gene-transfer agents [10, 16] and, usually overlooked, non-specifically stained organic and/or mineral particles [17, 18]. The latter might be important confounding factors in high-density ecosystems such as sediments and soils, where cell and VLP detachment from the organic-mineral matrix is required prior to epifluorescence-based quantification [19]. VMR values are particularly disparate in sediments and soils [19–21] and broadly vary depending on the protocol used [22]. Several observations suggest that the VLP counts proportionally diminish with increasing cell density [23, 24], although it has been hypothesized that lower VMR in cell-dense environments does not necessarily reflect low viral load but rather higher lysogenic prevalence [24]. However, while VMR seems to decline with increasing microbial cell densities, evidence for enhanced lysogeny remains ambiguous [25]. All these confounding factors challenge the accuracy of the existing VMR estimates and maintain uncertainty about the actual viral load supported by cellular organisms across ecosystems. Hence, more reliable approaches for obtaining this ratio and deriving ecologically relevant comparisons across biomes are needed.

Here, we develop a metagenome-based strategy to estimate VMR (mVMR) for DNA viruses (RNA viruses cannot be targeted by this approach) across all stages of their viral replication cycles (viral particles –VP–, as well as intracellular lytic and lysogenic viruses). We first benchmark this method on mock metagenomes with increasing VMR. Then, we estimate mVMRs from *de novo* generated aquatic-system metagenomes comprising coexisting viral particles (VPs) and cell fractions (50 kDa - 200 µm) and compare the obtained estimates with classical epifluorescence-derived estimates (fVMR). Finally, we estimate mVMR from 121 additional metagenomes across diverse ecosystem types, notably including high-density habitats. Our results show that mVMR are adequate proxies that can more realistically account for DNA virus:cell ratios in natural ecosystems than fVMR.

## Materials and Methods

### Metagenome-based counting of cells and viruses

To count cells, we used a set of 15 universal single-copy genes (USCGs; Table S1) highly conserved across the three cellular domains, including the reduced and often fast-evolving members of the Candidate Phyla Radiation (CPR, Patescibacteria) and the DPANN archaea. USCGs were identified via their PFAM motifs (Pfam-A database v3.1b2) [26] and searched in two datasets (Prokka-predicted genes and OrfM-predicted ORFs) with HMMER hmmsearch [27] v3.2.1 with the trust cut-off threshold. To assign relevant hits to taxa, they were analysed using Diamond [28] v0.9.25.126 against the NCBI non-redundant database formatted by Kaiju [29] v1.7.1: kaiju_db_nr_euk.faa, which contains all NCBI reference sequences for archaea, bacteria, eukaryotes and viruses. Best hits with more than 35% identity over at least 70% of query lengths were included in a best-hit table in kaiju-like output format using an *ad hoc* Perl script (mimicKaijuOutput.pl). They were then assigned to taxa using the kaiju script addTaxonName. Most genomes used to build the mock metagenomes had only one copy of the set of 15 USCGs (see below). Accordingly, the average of counts for the selected 15 USCGs in metagenomes represents an adequate proxy for individual cell counts. Five of the chosen ribosomal protein markers in the list of USCGs are known to have two copies in halophilic archaea and, accordingly, we divided by two haloarchaeal counts for them prior to averaging total counts. To estimate the number of viruses in metagenomes, we used as a proxy genes encoding major capsid proteins (MCPs). We used a total of 132 HMM profiles covering the MCPs of (almost) all known families of DNA viruses or viruses with intermediate DNA stages (ssDNA, dsDNA, RT-DNA) that infect bacteria, archaea and eukaryotes, including some relatively recently described viral groups, such as Yaraviridae [30]or Autolykiviridae [31] (but not the even more recently identified Mirusviruses [32], which might remain less well detected). For the analysis of the mock and planktonic metagenomes (see below), we specifically included HMM profiles of environmentally abundant marine viruses discovered by metagenomics (e.g., viruses of MGII Euryarchaeota, MGI Thaumarchaeota, marine Thermoplasmata, etc) and, to capture viruses of the rest of metagenomes, we built specific HMM profiles. For this, metagenome contigs assigned to viruses whose MCPs were not captured by existing HMM were used to build new HMM profiles. For reference, MCP sequences representing each family of DNA viruses, homologs were retrieved from the UniRef database, filtered to 30% sequence identity (3 iterations, inclusion value of E=1e-3) and aligned using hhblits [33], part of the hhsuite package 3.3.0 (Ref. [34]). The resulting alignments were converted to profile HMMs using hmmbuild included in the HMMER package v3.3.2 (Ref. [35]). Most genes retrieved with the MCP HMMs were functionally annotated as either capsid or hypothetical proteins and were assigned to viruses by our taxonomic assignment pipeline. However, several HMMs retrieved genes annotated as (diverse) encapsulins and simultaneously assigned to bacteria; these HMMs preferentially retrieving bacterial encapsulins were excluded from further analyses.

MCP HMM profiles that consistently retrieved genes assigned to cellular taxa and had clear cellular functions were discarded. A final set of 132 MCP HMMs (Table S2) was retained and used to search against all the Viral Ref Seq database (NCBI) using HMMER hmmsearch [27] v3.2.1 with an E value of 1e-11 threshold. In the case of genes attributed to different MCPs, the gene IDs were hashed and the MCP assignment with the lowest e-value was retained. Since multiple MCPs can occasionally be found in large viral genomes, we counted only one MCP per contig and considered that, under these constraints, 1 MCP represents 1 virus. To assess mVMR in mock metagenomes, nucleotide sequences of identified cellular USCGs and viral MCPs were indexed with Bowtie [36] v2.3.5.1 and the original mock metagenome reads mapped back onto them. Mapped reads were retrieved with Samtools [37] v1.9 and, for each gene, the Reads Per Kilobase (of gene sequence) per Million of mapped metagenome reads (RPKM) were calculated using *ad hoc* Perl scripts, allowing normalization for gene length and sample sequencing depth. Subsequently, we selected the USCG that recovered the most RPKMs per metagenome and the sum of MCP RPKMs to obtain normalised mVMRs. For an easier visualization of the proportion of viruses per microbial cell, we further normalised the viral RPKMs with respect to the number of cells (cell value = 1). Plots were generated with *ad hoc* R scripts using the ggplot2 package [38].

### Mock metagenomes

Ten mock metagenomes were built from a selection of 50 viral and 50 cellular genomes retrieved from the NCBI reference sequence database RefSeq (https://www.ncbi.nlm.nih.gov/) and GTDB (https://gtdb.ecogenomic.org/). Genomes were selected to partly mimic a simplified microbial community of marine plankton that included viruses, archaea, bacteria and eukaryotes (Table S3). To simulate the output of Illumina HiSeq sequencing, all genomic sequences were fragmented into 10 million reads of 100 bp using cMessi [39]. Eight mock metagenomes were built by varying the relative abundance of the viral vs. cellular reads in a range from 10 cells per viral genome to 125 viral genomes per microbial cell (V001M10 to V125M01; names reflect the virus:microbe ratio). The two additional metagenomes contained either none of the selected viral genomes (V000M01) or only viral sequences (V001M00; Fig. 1A). The generated reads were assembled with Metaspades [40] v3.13.1 with default parameters and k-mer iteration cycles of “21,25,31,35,41,45,51,55”. Assembly statistics are shown in Table S4. Gene annotation was performed on assembled contigs with Prokka [41] v1.14.5 in metagenome mode (contigs >200 bp). In parallel, to more exhaustively identify gene/protein domains, all ORFs were predicted with OrfM [42] (Table S5). Both annotation methods (Prokka and OrfM) produced very similar results (Fig. S1), albeit Prokka performed slightly better for high VMR values (Fig. 1B-C). We thus chose to display the contig-based Prokka annotation in subsequent analyses.

**Fig. 1.**
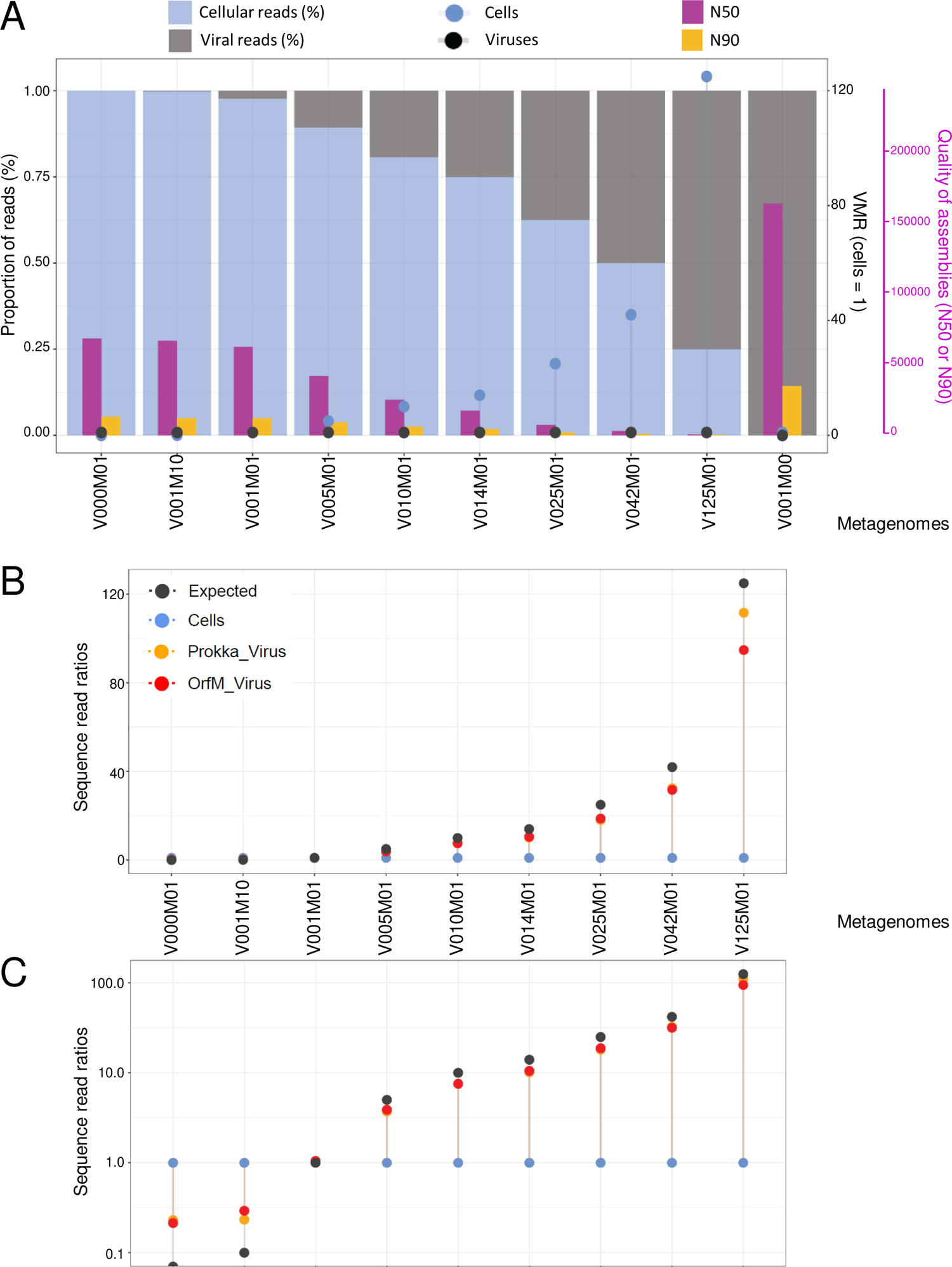
Estimated metagenome virus-microbe ratios (mVMR) across mock metagenomes with increasing VMR. **A**, Statistics of the different mock metagenomes generated with an increasing number of virus genomes (V000 to V125) as compared to the number of microbial cells (M00 – M01 – M10). **B**, Expected versus observed viral ratios versus cells (cells normalised to 1) in the different mock metagenomes as a function of the annotation method (OrfM vs. Prokka) used. Cells were counted using universal single copy genes (USCGs) and viruses using major capsid proteins (MCPs). **C**, Normalised counts (RPKM) of viruses per cell (normalised to 1) observed in mock metagenomes with our pipeline using OrfM and Prokka annotation strategies; a log10 y-axis scale is used for a better discrimination in metagenomes where viral genomes were either absent or in very low abundance.

### Water sampling, filtration, DNA purification and mixes for metagenome sequencing

Samples from the Mediterranean Sea were collected on 27/02/2019 at 20 m (MedS) and 40 m (MedD) depth in the water column (bottom depth 220 m) from a previously studied site off the Alicante coast (38°4’6.64’N, 0°13’55.18’W) [43]. The CTD (Seabird) data showed a typically winter-mixed water column, with homogeneous temperature (14.8°C), pH (8.4) and dissolved oxygen (8.5 ppm) throughout at least 60 m depth. Water was collected through a hose with a 200-µm Nitex mesh-capped end attached to the CTD and directly pumped onto a serial filtration system mounted with 20, 5 and 0.22 μm pore-size polycarbonate filters (Millipore). Biomass-containing filters were frozen on dry ice for transport to the laboratory prior to immediate DNA purification. The resulting <0.2 µm filtrate (200 l) was recovered in 50-l carboys and used for tangential filtration, which took place overnight on arrival at 4°C (cold room) using 6 Vivaflow200 filtration units (Sartorius) operating in two parallel series. The volume of concentrated viral fractions (>50 kDa) was further reduced using Amicon Ultra 15 ml Centrifugal Filters (Merck-Millipore) prior to phenol/chloroform extraction. Water samples (10 l) with increasing salt concentration (6%, 14%) were collected in carboys on 28/02/2019 from the Bras del Port solar saltern ponds: Sal6 (38°11’52.30’N, 0°36’13.15’W), Sal14 (38°12’4.74’N, 0°35’43.14’W). Brine samples were taken to the Alicante laboratory for immediate serial filtration (as above) and DNA purification. Freshwater (10 l) from a shallow lake in North-western France (Etang des Vallées; 48°41’23,0’’N, 01°54’59.2’’E) [44] was collected on 18/03/2019. Filtration was immediately carried out at the Orsay laboratory as above except we used filters of 100, 30, 5 and 0.22 µm pore-size, to avoid the many suspended plant material-derived particles in this shallow lake) In all cases, 4-ml aliquots of the different size-fraction filtrates and the final eluate (<50 kDa) were taken for flow cytometry and fixed overnight at 4°C with 0.5% glutaraldehyde then frozen in dry ice and stored at −80°C (Table S6). DNA from cellular fractions was purified from filters. These were cut into small pieces with a scalpel, added to 5 ml CTAB lysis buffer (SERVA) with 1 mg/ml lysozyme and incubated at 37°C for 1 h. After addition of proteinase K (0.2 mg/ml) and vortexing, the mix was incubated at 55°C for one additional hour prior to phenol/chloroform/isoamyl alcohol (25:24:1, pH 8) and chloroform/isoamyl alcohol extraction. Nucleic acids were precipitated overnight at −20°C with 1/10 volume of 3 M sodium acetate, pH 5.2, and 2.5 volumes of absolute ethanol. After centrifugation at 4°C, 13000 x g for 30 min, the pellet was washed with 70% ethanol, centrifuged again and speed-vac dried before resuspension in 10 mM Tris-HCl, pH 8.0. To purify DNA from viral concentrates, we incubated them with 20 mM EDTA, 50 ng/µl proteinase K and 0.2% SDS (final concentrations) for 1 h at 55°C and proceeded with the phenol/chloroform extraction as before. To recover the maximum of the aqueous phase, we used MaXtract High Density (Qiagen). DNA from the different cell fractions was fluorometrically quantified using Qubit and mixed proportionally according to the same amount of original water sample as indicated in Table S7. For all samples, we prepared a DNA mix “all” which included DNA from the viral and all cellular fractions below 200 µm (100 µm for EdV) and a mix “0.2v” which contained DNA from viral particles and cells smaller than 5 µm (prokaryotes and small protists). All these mixes were used for metagenome sequencing. As an internal control for our fractionation process, we also sequenced DNA from the viral fraction (metaviromes) from the Mediterranean plankton (Table S8).

### Flow cytometry

We quantified virus-like particles (VLP) and cells (up to ∼10 µm) by flow cytometry (FCM) in the glutaraldehyde-fixed samples spanning different salinities and size fractions described above (Table S6). We used a CytoFLEX S (Beckman Coulter) equipped with two active diode lasers (488 nm and 561 nm) following well-established protocols [45, 46]. The flow cytometer settings and gains were, for VLP detection, forward scatter (FSC) 500, side scatter (SSC) 500 and green fluorescence (FITC) 400 with a threshold set to FITC 260; and, for cell detection, FSC 250, SSC 700 and FITC 70 with a threshold set to FITC 1000. The sheath fluid consisted of 0.1 µm-filtered Milli-Q water. Multifluorescent beads of 0.2 and 1µm (Polyscience) were used as size reference for VLP and cells. Samples were thawed on ice and diluted up to 1000 times for VLP counts and 10 times for cell counts with 0.02 µm-filtered autoclaved Milli-Q water, TE buffer (Sigma) or seawater. VLP and cells were stained with SYBRGreen I (Sigma) for 10 min in the dark at 80°C and let cool down for 5 min at room temperature [46]. Background noise was systematically checked on blanks (Fig. S2). Samples were recorded with an event rate per second of 100-1,000 (VLP) and <300 (cells). For VLP, we defined three gates (V1, V2, V3) on a FITC-A vs SSC-A plot (Fig. S2). The VLP signal intensity was determined on marine plankton controls, with gates set to exclude background based on FITC signal. Cells were stained with SYBRGreen I for 10 min at room temperature in the dark. For cell counts, we established two gates, HDNA and LDNA, on a FITC-A *vs* SSC-A plot, as previously described [45]. Sample counts were analysed with CytExpert and Kaluza softwares (Beckman Coulter). VLP and cell counts were obtained by correcting the measured total counts for noise (typically having the lowest green fluorescence) using blanks. VLP and cell abundances were respectively expressed as VLP.ml^-1^or Cells.ml^-1^ (Table S6).

### DNA purification of sediment and microbial mats, sequencing and selection of natural metagenomes

We also included in this analysis metagenomes of freshwater sediments and microbial mats from previous and ongoing studies in our laboratory (Table S8). DNA from Lake Baikal sediment samples collected in July 2017 was purified using the Power Soil DNA purification kit (Qiagen, Germany) [47]. DNA from microbial mat samples collected from the Salada de Chiprana (Spain, December 2013), lakes Bezymyannoe and Reid (Antarctica, January 2017) and several hot springs around Lake Baikal (Southern Siberia, July 2017), fixed *in situ* in ethanol as previously described [48], was purified using the Power Biofilm DNA purification kit (Qiagen, Germany). DNA was quantified using Qubit. Sequencing (Illumina HiSeq, paired-end, 2×125 bp) was performed by Eurofins Genomics (Ebersberg, Germany). In total, we generated 12 metagenomes from planktonic fractions coming from increasingly saline waters (freshwater, marine and solar saltern ponds at 6-14% salt), 8 metagenomes from freshwater sediment and 23 metagenomes of diverse microbial mats. To these, we added 13 metagenomes of other microbial mats and 7 from microbialites (18 of which were previously generated in our laboratory), 26 metagenomes from lake sediments, 6 marine sediment metagenomes, 16 soil metagenomes and 30 animal-associated microbiomes from diverse organs and animals (honey bees, frogs, humans), all of which were sequenced with the same technique (Illumina). In total, we included 133 metagenomes in this study (2 of them were control Mediterranean metaviromes), 40 metagenomes generated in our laboratory and 93 metagenomes from other published studies [48–61] (an extended description, GenBank accession numbers and statistics are provided in Table S8).

### Sequence analysis, annotation and taxonomic assignation of natural metagenomes

We treated the 133 metagenomic datasets with exactly the same bioinformatics pipeline starting from raw Illumina sequences. Read quality was verified with FastQC [62] v0.11.8 and cleaned with Trimmomatic [63] v0.39, adjusting the parameters as needed (usually LEADING:3 TRAILING:3 MAXINFO:30:0.8 MINLEN:36) and eliminating the Illumina adapters if any. Clean reads were assembled with Metaspades [40] v3.13.1 with default parameters and k-mer iteration cycles of “21,25,31,35,41,45,51,55”. A few complex metagenomes that failed to assemble in our server were assembled with Megahit [64] v1.2.6 with default parameters except for a minimum contig length of 200 bp and starting kmer sizes of 23 to 93 with an increasing step of 10 (Table S8). Gene annotation was performed with Prokka [41] v1.14.5 in metagenome mode (contigs >200 bp). We assigned coding sequences to PFAMs (Pfam-A database v3.1b2) with HMMER hmmsearch [27] v3.2.1 with the trust cut-off threshold. As PhiX174 genomic DNA is sometimes added during sequencing as internal control, we downloaded the reference phage PhiX174 genome and BLASTed the 11 encoded proteins against the set of significant MCPs of each metagenome. Hits to PhiX174’s MCP (NP_040711.1) with 100% identity over a 90% coverage were excluded from the analysis. Taxonomy was assigned to each gene in metagenomes as described above for the mock metagenomes. Taxon-assigned genes were then classified using an *ad hoc* Perl script (assignTaxa.pl) into the following major groups, further classified in three major lifestyle categories: i) CPR bacteria (Patescibacteria) and DPANN archaea (Table S9), considered as cellular episymbionts/parasites strictly dependent on free-living organisms; ii) “Other Archaea”, “Other Bacteria” and “Eukaryotes”, classed as free-living (FL), and iii) viruses. An *ad hoc* Perl script (tax2PFAM.pl) linked each gene’s taxonomy to its Pfam assignment. Metagenome-inferred proteins with >35% amino acid identity over >70% sequence length to viral genome-encoded proteins were assigned to the respective family, host and virus group. Viral taxonomy and Baltimore classification were determined with the VPF-Class tool [65] (Table S10). MCP-containing contigs were used as an input for VPF-Class; we attributed a classification when values were over a membership ratio score of 0.75. The relative abundance of MCPs from these contigs in metagenomes were plotted using the R ggplot2 package visualizing the different viral families and Baltimore classification.

### Diversity indices

To calculate diversity estimates, we selected the most abundant USCG per group of metagenome (Table S11). Amino acid sequences for each ribosomal protein were retrieved with *ad hoc* Perl scripts for each metagenome group and clustered with cd-hit [66] at 99% identity (parameters -c 0.99, -n 5, -d 0, -T 16, -M 0). These clusters were considered operational taxonomic units (OTUs) and were assigned to CPR, DPANN, (non-DPANN) archaea, (non-CPR) bacteria and eukaryotes. For viruses, we used the whole set of viral MCPs under the premise that most viral genomes have only one MCP (see above). We obtained a total of 8,610 cellular clusters and 28,319 MCP clusters using Prokka annotation. Abundance matrices of these clusters were built with an *ad hoc* Perl script and several diversity indices (alpha diversity, Shannon entropy, Pielou’s evenness and Simpson’s dominance) were calculated with an *ad hoc* R script using the Vegan package [67]. Indices for all metagenomes were plotted with ggplot2. To compare if the difference between the viral and cellular diversity was significant, a Wilcoxon test was applied to the median of the alpha diversity per ecosystem, using an *ad hoc* R script with the ggpubr library. Visualization of the statistical test was represented as a box plot with the same R script and the ggplot2 library. Principal component analyses (PCA) of different diversity indices as a function of metagenomes were carried out with an *ad hoc* R script, and visualized using the ggplot2 package.

## Results

### Benchmarking mVMR in mock metagenomes

To develop a metagenome-based VMR estimate, we used universal single-copy genes (USCGs) and viral major capsid protein (MCP) genes, usually present in single copy in viral genomes, as proxies for cells and DNA viruses, respectively. We selected 15 ribosomal proteins that are well-represented across the three domains of life as USCGs (Table S1). To limit differential representation of individual ribosomal protein genes, we averaged values for the 15 USCGs as a proxy for cell counts under the assumption that in most biomes cells contain one genome copy. Relatively rare cases of polyploidy have been described, e.g. in giant bacteria [68, 69] and haloarchaea [70], and two chromosome copies usually occur in some archaea and actively transcribing bacteria [71]. However, in the wild, especially in energy-limited ecosystems, microbial cells are likely to exist in low average metabolic states and, consequently, limited ploidy. Should average ploidy be higher than one, this proxy might slightly overestimate cell numbers. Conversely, some markers do not capture well episymbiotic prokaryotes with reduced and divergent genomes, such as Candidate Phyla Radiation (CPR or Patescibacteria) bacteria and DPANN archaea [72, 73], which might partly compensate for the ploidy excess in some cells. To count viruses, we used 132 hidden Markov models (HMMs) that recognize MCPs from virtually all known families of DNA viruses infecting bacteria, archaea and eukaryotes. In addition, to avoid missing MCPs from potential unknown viruses present in the studied natural metagenomes (see below), we designed specific HMMs for viral genomes assembled from them (Table S2; see Methods).

We then generated 10 mock metagenomes with increasing VMRs, ranging from ratios of 10 cells per viral genome (V001M10; the number following V and M indicates the ratio of viral to microbial cell genomes per mock metagenome) to 125 viral genomes per microbial cell (V125M01). We also included one metagenome with only cellular genomes (V000M01) and one with only viral genomes, i.e. a metavirome (V001M00). These mock metagenomes partly mimicked marine planktonic communities, and included genomes of archaea, bacteria, eukaryotes and viruses (Table S3).

The assemblies in the mock metagenomes with the highest VMR (V042M01 and V125M01) were highly fragmented, exhibiting lower quality (Fig. 1A; Table S4). To cope with potential gene detection problems in these assemblies, we applied two annotation strategies, one based on predicted gene annotation (Prokka [41]) and another based on the annotation of all possible open reading frames (OrfM [42]) allowing for the detection of individual protein domains even if genes were fragmented. Even if many more ORFs than genes were detected per mock metagenome (Fig. S1A), the amounts and relative proportions of USCGs per metagenome were remarkably similar between the two methods except for those with the highest VMR, where OrfM performed quantitatively better (Fig. S1B). The relative proportions of high-rank cellular taxa obtained with the two annotation strategies were closely similar (Fig. S1C-D) and in agreement with the expected taxonomic assignment (Table S5). To estimate mVMR, we searched for USCGs and MCPs in the assembled contigs from these 10 metagenomes using the respective protein family HMMs (see Methods) and, after mapping the original reads onto the identified contigs, obtained normalised USCG and MCP numbers by counting the reads per kilobase of gene sequence per million of mapped metagenome reads (RPKMs). The inferred mVMR values were also similar for the two annotation strategies, and close to the expected VMR although, at very high VMR (V125M001), OrfM yielded slightly lower mVMR values than Prokka (Fig.1B). As expected, we could not detect USCGs in genes/ORFs predicted from the virus-only mock metavirome (V001M00). However, we did identify several MCPs in the metagenome containing only cellular genomes (V000M001). These were annotated as cellular virus-related or hypothetical proteins/domains by Prokka and corresponded to proviruses present in the included cellular genomes (Table S3). Accordingly, it was possible to differentiate lysogenic viruses from VPs and intracellular lytic viruses in these mock metagenomes (Fig.1C; Fig. S1C-D). The presence of proviruses likely explains the observation that, at very low VMR (V000M001-V001M10), viral counts appeared slightly overestimated (Fig.1C).

### Estimation of mVMR in aquatic metagenomes

After benchmarking mVMR in mock metagenomes, we first sought to estimate mVMRs for aquatic ecosystems and compare the resulting values to classical epifluorescence-derived estimates (fVMR). However, published planktonic metagenomes are not suitable for the estimation of total mVMRs due to an inherent problem resulting from the fact that DNA is purified from the planktonic biomass retained on different pore-size filters upon filtration. Most of the available aquatic metagenomes contain the prokaryotic/picoeukaryotic fraction (0.2 to 3-5 µm cell size) and, more rarely, larger-size fractions where many protists are retained (>5 and up to 100-200 µm). Metagenomes from smaller plankton fractions containing VPs (“metaviromes”) are obtained after tangential filtration of <0.2 µm filtrates to retain particles larger than 50-100 kDa. However, many VPs can be also retained in pre-filtration steps and end up in “cellular” fractions, for instance retained on membranes in-between pores and, especially, trapped among cells as filters saturate with biomass. A corollary is that cellular metagenomes and metaviromes are usually generated from different water volumes/samples, after imperfect particle fractionation steps, sequenced from different libraries and analysed separately, such that mVMR cannot be directly and reliably estimated. To overcome this problem, we generated integral microbial plankton metagenomes, including genomes of both cells and viruses (covering both extracellular and intracellular phases of the virus replication cycle), from the same water mass of selected aquatic ecosystems (Fig. 2). They included a freshwater shallow lake in Northern France (Etang des Vallées, EdV), the Western Mediterranean water column (sampled at 20 and 40 m depth) and two ponds of increasing salinity (6% and 14% salt) from a Mediterranean solar saltern (Table S8). To obtain more representative VMR, we sampled these ecosystems in late winter (see Methods), avoiding massive blooming periods when richness and evenness decrease [74].

**Fig. 2.**
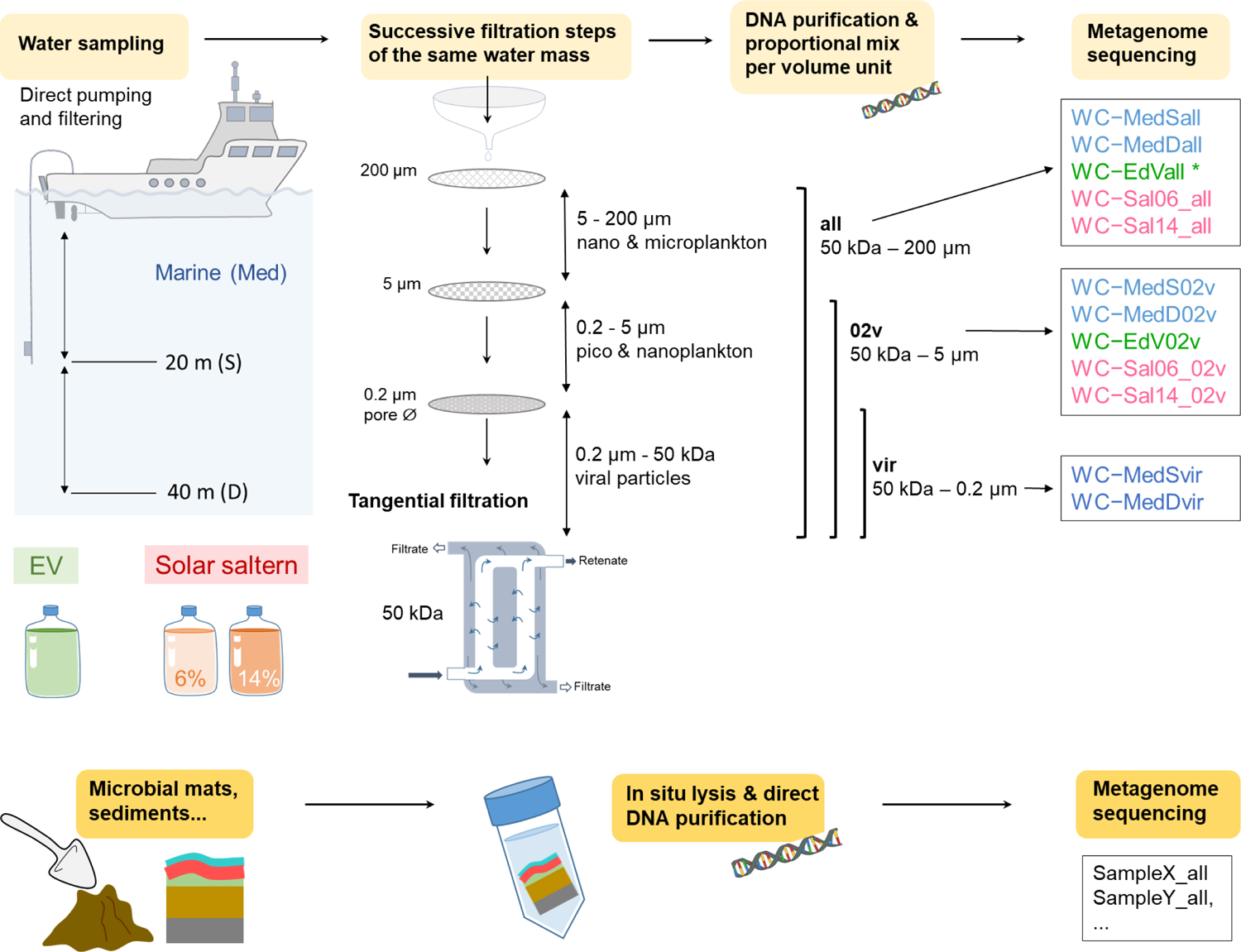
Strategies used to generate metagenomes in this study. Metagenomes from different planktonic fractions were obtained after sequential filtration of the same water mass, purification of DNA for each size fraction and proportional mixing per volume unit. For simplification, the intermediate filter of 20-30 µm pore-diameter size is not shown. Two metaviromes (particles <0.2 µm) were sequenced for control. Metagenomes of non-aquatic samples were generated after the in situ cell lysis and direct DNA purification prior to sequencing. EV, Etang des Vallées (freshwater system). * The metagenome WC-EdVall corresponds to the size fraction 50 kDa – 100 µm.

To generate metagenomes of coexisting viral and microbial cell plankton, we carried out serial filtration steps from the same initial water mass through filters of 200 µm pore-diameter to exclude metazoans larger than that size and then 20, 5 and 0.2 µm, retaining cells of different sizes. Tangential filtration was applied to the resulting <0.2 µm-filtrate to concentrate viral particles (>50 kDa) (Fig. 2). We retained aliquots of all sequential filtrates for fluorescent cytometry (FCM) counts of labelled cells and VLPs, and the final eluate (<50 kDa) as a blank (Table S6 and Fig. S2). FCM counts of cells in all fractions >0.2 µm were consistently similar for each sample type, indicating that the vast majority of cells were prokaryotic. Cell abundances were higher in saltern ponds (5.2×10^6^ to 1.0×10^7^ cells.ml^-1^) and freshwater samples (4.3 to 4.7×10^6^ cells.ml^-1^) compared to marine samples, for which the counts were about an order of magnitude lower (4.5 to ×10^5^ cells.ml^-1^) (Fig. 3A; Table S6). VLP counts were also consistent in large-size fractions of each sample and diminished in the tangential-filtration concentrate. This observation indicates that many VLPs were retained in larger fractions due to progressive filter saturation with biomass and highlights the importance of combining different size fractions of the same filtration series for metagenome-based abundance estimates. VLP abundances appeared higher in saline environments (5.6×10^7^ and 4.1×10^7^ VLP.ml^-1^ for 6% and 14% salt ponds, respectively). Although this could reflect the actual abundance of viruses, it is also possible that salt nanoprecipitates forming at increasing salinities get unspecifically fluorescently stained [18]. Estimated fVMRs ranged from ∼3 to 15, with lower values for freshwater (∼4), 40 m-deep Mediterranean (∼5) and 14%-salt pond (∼3) and higher for 20 m-deep Mediterranean (∼14) and 6% (∼10) saltern ponds (Fig.3A and D; Table S6).

**Fig. 3.**
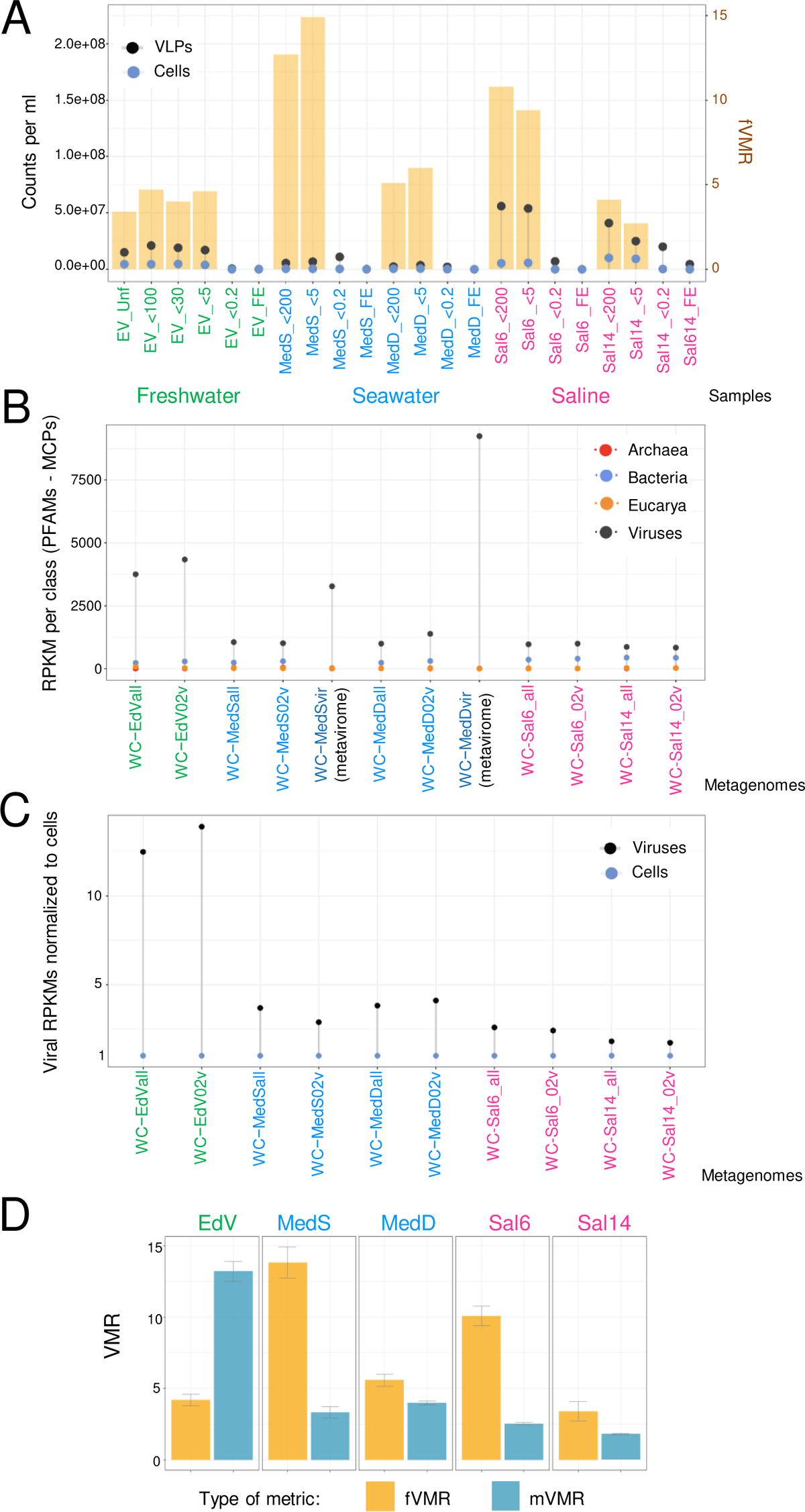
Epifluorescence- and normalised metagenome-based counts and ratios of cells and viruses in aquatic ecosystems. **A**, Flow-cytometry counts of cells and viral-like particles (VLPs) and estimated virus-to-microbe ratios (fVMR; histogram bars) from the same aquatic samples used to determine mVMR: EV, Etang des Vallées; MedS and MedD, 20 m and 40 m-deep Mediterranean plankton, respectively; Sal06 and Sal14, saltern ponds of 6 and 14% salinity (w/v), respectively. Sample names also indicate the plankton size fraction considered (Unf, all size fractions; <200, <100, <30 and <5, fractions below those numbers in µm; <0.2, fraction between 50 kDa and 0.2 µm; FE, final eluate <50 kDa). **B**, Normalised counts (RPKM) of viruses (MCPs) and cells (USCGs) from the three cellular domains metagenomes from the same original aquatic samples as in (A). Metagenomes correspond to size plankton fractions indicated by their suffixes: -all, fraction between 50 kDa and 200 µm (100 µm for EV); 02v, fraction between 50 kDa and 5 µm. Two metaviromes (fraction 50 kDa-0.2 µm) were sequenced for Mediterranean (MedS, MedD) plankton samples as internal control. **C**, Metagenome-based normalised ratios of viruses per cell (bacteria + archaea + eukaryotes). RPKM, reads per kilo base per million mapped metagenome reads. **D**, Comparison of observed fMVR and mVMR for the same samples.

We purified DNA from the biomass retained on all filters used during serial filtration as well as from tangential-filtration concentrates, keeping track of the corresponding water volumes. DNA samples obtained from different fractions were then proportionally mixed for a given original water volume (Table S7). We sequenced metagenomes from two plankton fractions including viral particles and cells up to i) 5 µm, i.e. prokaryotes and pico-nanoeukaryotes (50 kDa - 5 µm; “02v” fraction), and ii) 200 µm (100 µm for EdV), including unicellular prokaryotes and pico-nano-microeukaryotes (50 kDa - 200 µm; “all” fraction) (Fig. 2). In this way, co-occurring viral and cellular DNA was sequenced from the same library, avoiding technical biases. As eukaryotes represented a minor fraction in these aquatic systems, the two metagenomes were expected to behave as quasi-replicates for mVMR inference. In addition, we sequenced two “metavirome” fractions (50 kDa - 0.2 µm; “vir” fraction) for the marine samples as internal controls (Table S8). Normalised numbers of USCGs were generally congruent in our aquatic metagenomes, although with slight among-marker differences (Fig. S3). In all our planktonic metagenomes, normalised viral counts (MCPs) were higher than USCG-based cell counts (Fig. 3B). Most cell counts were bacterial; as expected, eukaryotic counts were comparatively low (Fig. 3B).

Estimates of mVMR in our aquatic ecosystems varied between 1.3 and 10.5, with marine samples exhibiting values of 3-4 (Fig. 3C-D; Table S8). mVMRs were lower than fVMR in marine and saline samples (Fig. 3D), although the two values were comparable in the case of MedD (∼4-6) and Sal14 (∼2-3). By contrast, in the case of freshwater EdV samples, mVMR values were 3 times higher than fVMRs. This difference seems to suggest that, in addition to VPs, intracellular lytic viruses and/or proviruses were highly prevalent in the plankton of this highly diverse shallow lake [75].

### mVMR across ecosystems

We subsequently calculated mVMRs in additional metagenomes (for a total of 131, leaving aside the two metaviromes; Table S8) covering a wide biome diversity including marine and lake sediments, soils, microbial mats growing in various conditions (halophilic to thermophilic), microbialites and several types of metazoan-associated microbiomes. Many of those metagenomes were retrieved from public databases but 40 of them were generated in our laboratory, notably from poorly studied ecosystems (Table S8). To avoid potential confounding factors derived from sequencing techniques and bioinformatic analysis, all chosen metagenomes were sequenced using the same technology and treated uniformly starting from raw reads. Although normalised viral MCPs tended to dominate over cellular USCGs across the 131 studied metagenomes, cellular markers dominated in about a third of them and, in several metagenomes, viral and cellular counts were comparable (Fig. 4; Fig. S4-S5).

**Fig. 4.**
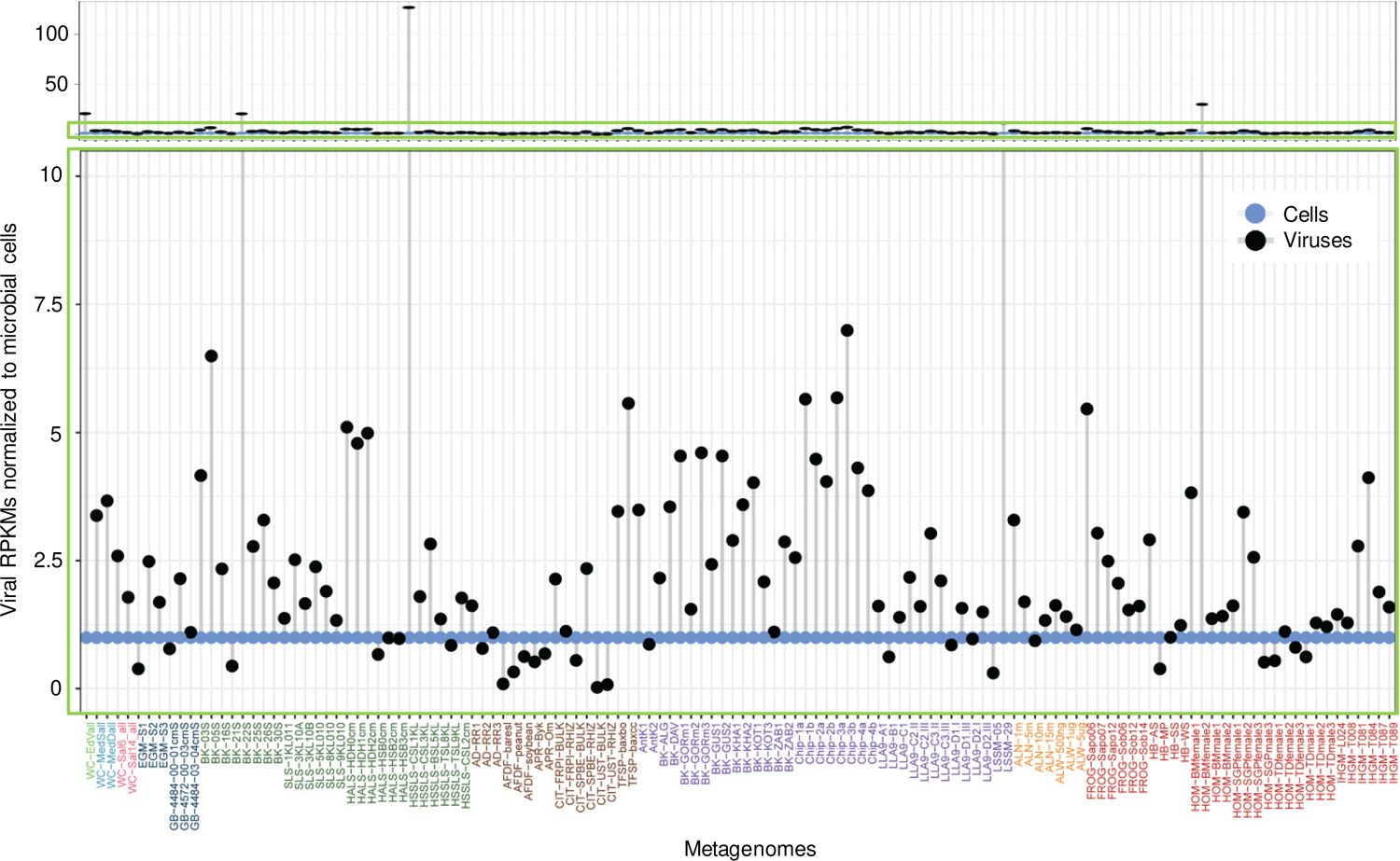
Metagenome-based normalised ratios of viruses per cell across ecosystem types. Viruses and cells were counted using, respectively, MCPs and USCGs as proxies; counts were normalised to cell = 1. The lower panel corresponds to a zoom of the area shown within a green rectangle in the upper panel. All the metagenomes were analysed using the same pipeline from raw reads. Metagenome labels in the x-axis are coloured according to their ecosystem type as indicated. RPKM, reads per kilo base per million mapped metagenome reads.

At finer taxonomic scales, the diversity and relative proportions of cellular organisms and viruses were overall congruent with those reported for the different types of ecosystems and the specific studied metagenomes (Table S8). Bacterial taxa were largely dominant across biomes, although archaea and microbial eukaryotes were also present (Fig. 5A; Fig. S5). Archaea were abundant in marine sediments [49], several microbial mats [48] and the surface Mediterranean waters, the latter dominated by Marine Group II archaea (Fig. 5A). Diverse Alpha- and Gammaproteobacteria were highly abundant in marine plankton but also in microbialites and, to a lesser extent, microbial mats. Cyanobacteria were also present in considerable proportions in these ecosystems. Actinobacteria and Acidobacteria were dominant in soils. Some actinobacterial taxa were also abundant in the freshwater system [75]. Microbial eukaryotes consistently occurred in most ecosystems, albeit in minor proportions. They were more abundant in plankton samples and microbialites; diverse microalgae, notably diatoms and chlorophytes dominated the eukaryotic component (Fig. 5A). The abundances of CPR and DPANN (the latter included in “other archaea” in Fig. 5A) were generally relatively low (albeit reaching ∼10% in some samples) and did not outnumber viruses in the same ecosystems, suggesting that viral parasitism is generally more prevalent than prokaryotic parasitism.

**Fig. 5.**
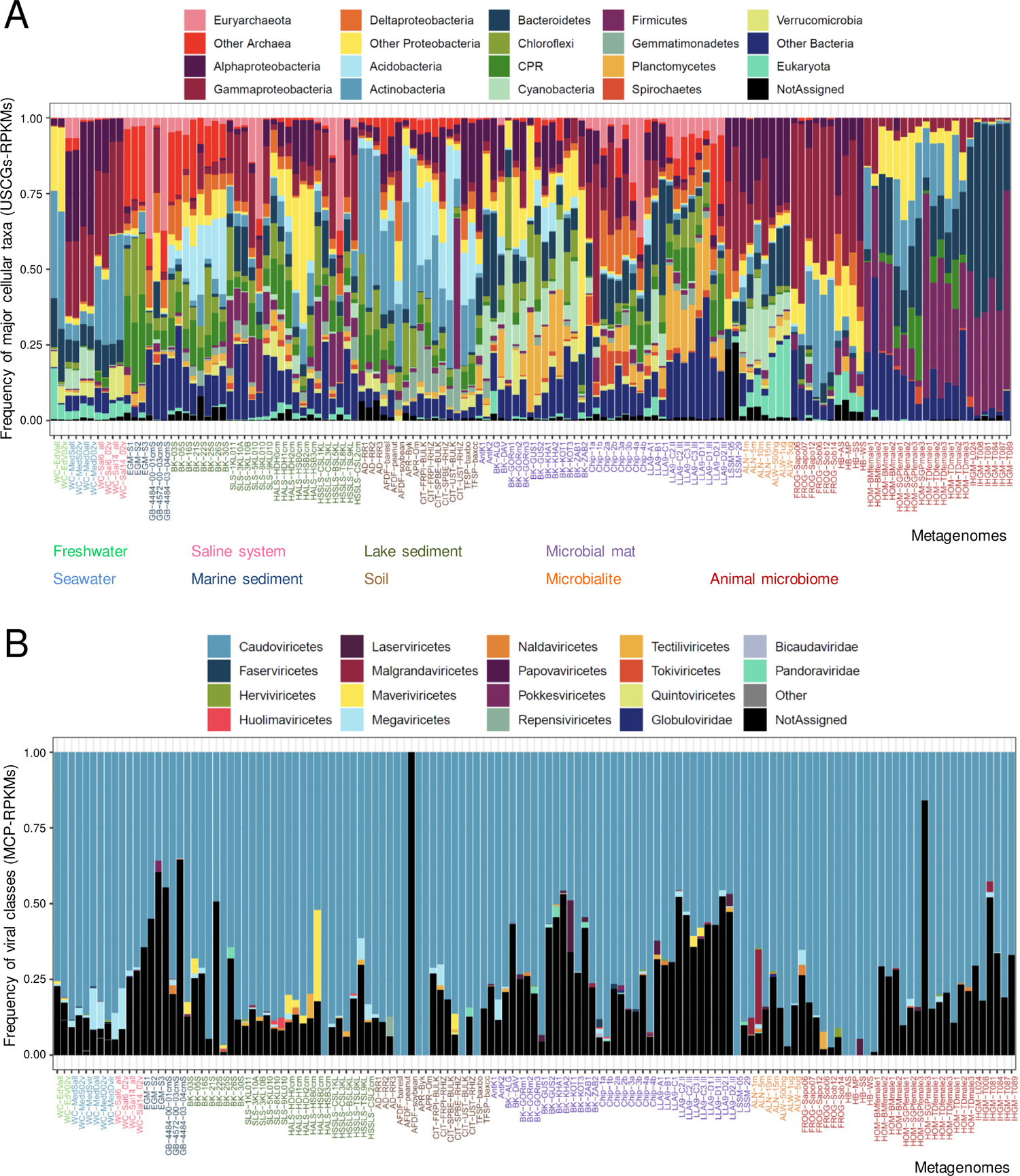
Frequency of major cellular and viral taxa identified in all studied metagenomes based on the phylogenetic assignment of USCG and MCPs. Metagenome names are coloured depending on their ecosystem type as indicated.

Viruses were also highly diverse although, as expected, dsDNA viruses largely dominated the studied microbial ecosystems; ssDNA viruses were also detected (Fig. S6; Table S10). In agreement with the pronounced dominance of bacteria as potential hosts in most biomes, the dominant identified viral families belonged to the class *Caudoviricetes* (tailed bacteriophages and related archaeal viruses [76], containing the HK97-fold MCP), which collectively represent ∼74% (n=8281) of all identified viral contigs across ecosystems. The bacteriophage family *Microviridae*, with ssDNA genomes, was represented in some ecosystems. Among eukaryotic viruses, the most represented were members of the *Phycodnaviridae*, which typically parasitize algae, and *Mimiviridae*, which infect diverse protists (e.g. amoeba) [77] (Fig. 5B; Table S10).

In summary, although the inferred mVMR values were on average greater than 1 (global average 1.96, i.e. ∼2 viruses per cell), mVMR values were lower than 1 in approximately one third of the metagenomes (Fig. 4). In only 5 out of the 131 metagenomes, viruses exceeded cells by more than one order of magnitude. The rest of the metagenomes displayed mVMR values between 1 and ∼7. In terms of grand ecosystem types, mVMR were the highest in aquatic ecosystems (mean 4.0; median 3.0) and the lowest in animal-associated microbiomes (mean 1.3; median 0.8; Fig. 6A), consistent with the low recent VMR estimates in the human gut [78]. Microbial mats and soils/sediments exhibited intermediate average values (respective means, 1.5 and 3.1, and medians, 1.1, 1.0), the latter displaying a wider value distribution with a higher variance, in agreement with previous observations [19–21].

**Fig. 6.**
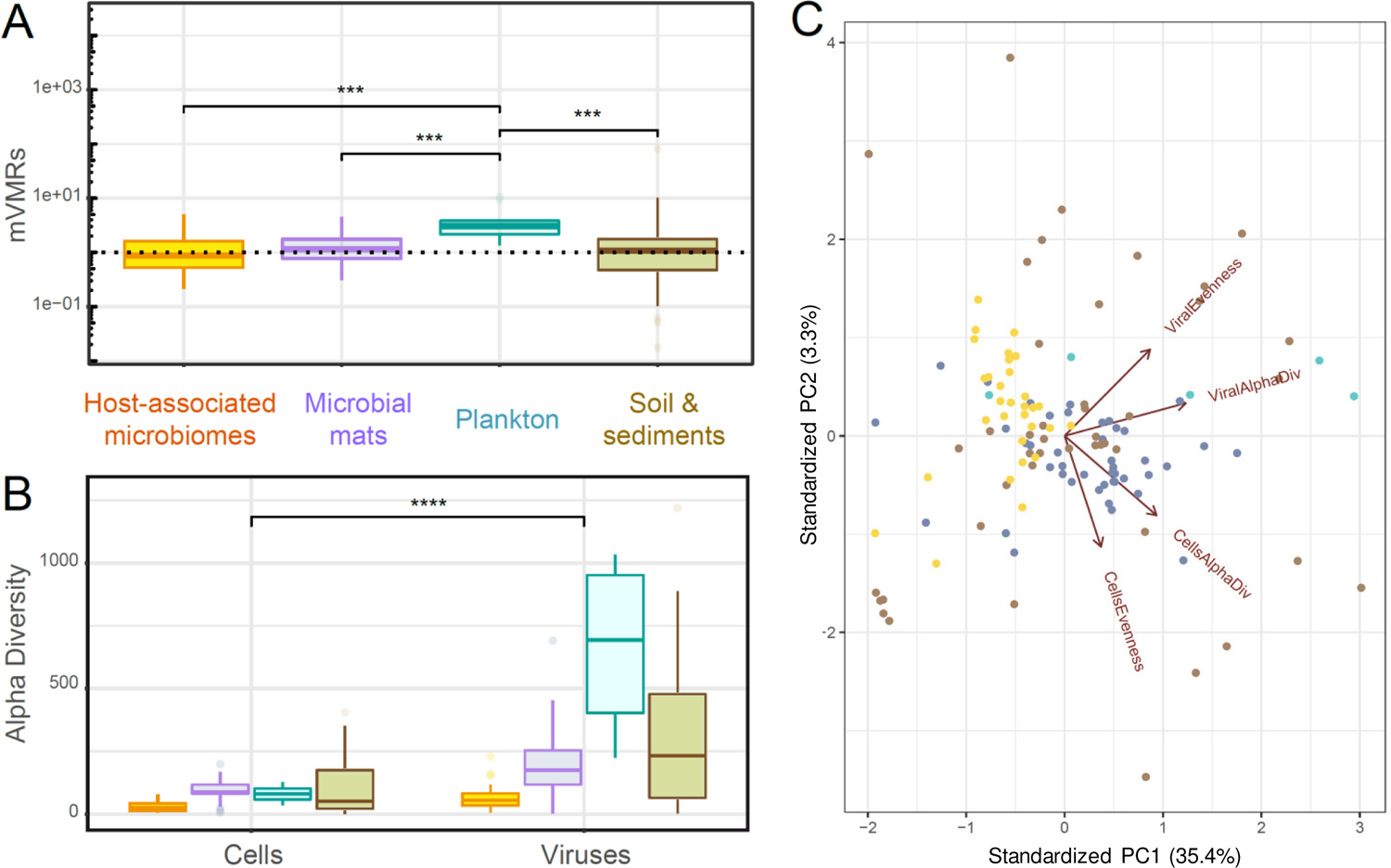
mVMR and diversity parameters of cells and viruses per grand ecosystem type. **A**, Boxplot showing the distribution of mVMR values; only significant differences are shown (*p*<0.001). **B**, Boxplot showing the alpha diversity of cells and viruses; diversity differences between cells and viruses were significant (*p*<0.0001). **C**, Principal component analysis of different metagenomes maximizing the variance explained by viruses (MCPs counts in RPKMs) and microbial cells (USCG counts in RPKMs). Metagenomes are coloured as a function of grand ecosystem type.

To look for potential determinants explaining the distribution of viruses and cells across biomes, we carried out principal component analyses (PCA) of the studied metagenomes using diversity-related parameters, namely richness, alpha-diversity and evenness indices for cells and viruses (Fig. S7). As expected, cell alpha-diversity positively correlated with microbial mats and soils, being lower in host-related microbiomes, whereas viral diversity best correlated with marine and freshwater plankton. Cell evenness mostly correlated with some soil and microbialite samples, whereas viral evenness correlated with freshwater sediment samples (Fig. 6 for grand ecosystem types; Fig. S8 for more detailed ecosystems). Globally, alpha-diversity values estimated for cells and viruses were significantly different among specific (Fig. S8B) and grand ecosystem (Fig. 6B) types. The alpha-diversity of viral plankton was the highest followed, albeit at a distance, by sediments, both freshwater and marine (Fig. 6B; Fig. S8B). This trend, along with higher viral abundances, is consistent with the notion that water masses constitute a reservoir of dispersing viral particles. By contrast, highly structured and heterogeneous environments are less prone to host a wide and abundant diversity of dispersing forms because, in the absence of their specific hosts, these particles can be easily used as food or otherwise degrade. Freshwater and marine sediments would appear as intermediates, receiving a constant input of sedimenting VPs, which can then enter a decay and degradation process associated to local prokaryotic communities.

## Discussion

In this work, we developed and benchmarked on mock metagenome datasets a metagenome-based VMR metric based on virus and cell markers that is applicable throughout microbial ecosystems in a comparative manner. Our mVMR metric is *a priori* more accurate for the inference of actual (DNA) virus:microbe ratios because, in comparison with fVMR values based on fluorescently labelled VLPs only, it accounts also for the actively infecting, intracellular lytic viruses as well as lysogenic forms. Our approach is thus more viral-inclusive and free from potential biases affecting fVMR [10, 12–14, 16–18]. In addition to identifying specifically stained vesicles and capsids containing non-viral DNA (transducing phages or gene transfer agents), fluorescence-based methods are affected by biases stemming from the differential treatment of contrasting sample types (water, soil, sediment, mucilaginous mats) and can include non-specifically stained organic and/or mineral particles from complex ecosystems. These artefacts may not only affect VLP counts, but possibly also cell counts, casting doubt on total cell estimates based on these methods in sediments, soils or the subsurface and, hence, global virus and cell counts [9, 10]. Furthermore, because a sizable fraction of viral particles is defective [79–81], the real number of infective viruses derived from VLP counts remains uncertain.

Our approach is not fully devoid of potential biases that might result in slight relative over- or underestimates of cells and/or viruses, such as differences in cell ploidy or divergent viral and cellular individual gene markers. However, as metagenome sequencing and the exploration of cellular and viral diversity across ecosystem types improve, detection problems linked to sequence divergence should be gradually solved. Obviously, being metagenome-based, our approach does not allow for the quantification of RNA viruses and, for reverse-transcribing viruses and single-stranded DNA viruses, only the replicative DNA intermediates can be identified. Notably, epifluorescence-based approaches are equally unsuited for enumeration of viruses with small RNA and ssDNA genomes due to lack of sensitivity [82]. Thus, the numbers of their virus particles will be slightly underestimated using both the metagenomic approach described herein and the traditional fluorescence-based counting. Unfortunately, a similar approach based on metatranscriptomic counts is not feasible to infer VMRs for RNA viruses because USCGs can be expressed at drastically different levels in different cells, preventing them to be used as markers for individuals. Like DNA viromes uncovered by metagenomic studies [7, 83], the diversity of the global RNA virome is rapidly expanding thanks to metatranscriptomic analyses [84–86]. Nonetheless, dsDNA viruses dominate the prokaryotic virome [77] and, although several novel families of RNA viruses seem to infect bacteria [84, 85], bacterial viruses jointly comprise only 3.4% of the RNA viral OTUs [84]. Therefore, given that most microbial ecosystems are overwhelmingly dominated by prokaryotes, mVMR estimates based on gene-marker proxies for individual viruses and cells, even if in need of further refinement (especially with progressively expanding databases), are likely to be the best current approximation of the real VMR, at least at the order-of-magnitude level.

This work shows that mVMRs are higher in planktonic systems approaching, albeit not necessarily reaching, the values of ∼10:1 that are traditionally assumed to be the minimal values in current (fVMR-based) estimates (Fig. 6). Water masses therefore seem to represent reservoirs of viral particles that can more easily escape from cellular predation, whereby nucleic acids serve as an excellent source of P and N, even if the uptake of specific lytic viral nucleic acids comes at a cost [87]. From this perspective, aerosols likely are another huge reservoir of VPs. By contrast, in higher-density biomes, mVMR values appear substantially lower, consistent with higher viral dispersal limitation and degradation rates, as well as with the idea that VMR declines as a function of microbial cell densities [23]. Even if mVMRs are highly variable, especially in sediments and soils, mVMR estimates across non-aquatic ecosystems barely reach 2, being especially low in cell-dense multicellular host-associated microbiomes (<1). Overall, our mVMR estimates across ecosystems question the currently widely held assumption that viruses outnumber cells at least by one order of magnitude. Even if ocean waters occupy an important fraction of Earth’s volume, continental biomes together with the deep subsurface undoubtedly host most of the planet’s biomass and cells [88, 89].

Accordingly, although viruses likely are the most diverse biological entities on Earth [6, 7], notably because most cellular organisms can be infected by different viruses, they might quantitatively be almost at equal footing with cells. In addition, deciphering the proportion of actively infecting viruses should provide more useful information about the maximum load of viral entities that microbial communities can support before ecosystems collapse. Future exploration of these poorly studied parameters across ecosystems and along temporal series should help understanding the dynamics of viral-cell interactions in the biosphere.

## Acknowledgements

We thank José Manuel Haro Moreno for help during marine and solar saltern sampling in Alicante, Paola Bertolino for help during sampling in several other ecosystems, and Antonio Camacho y Antonio Picazo for CTD measurements during marine plankton sampling. We thank the Salinas Bras del Port for sampling permission and the UNICELL platform for flow cytometry analyses. We acknowledge support from the Moore-Simons Project on the Origin of the Eukaryotic Cell through Grant GBMF9739 (P.L.-G.) and the European Research Council Advanced Grants FP7 No. 322669 (P.L.-G.) and H2020 No. 787904 (D.M.).

## Data and code availability

Metagenome Illumina sequences generated in our laboratory and not yet published have been deposited in GenBank (National Center for Biotechnology Information) Short Read Archive with BioProject number PRJNA756245. Accession numbers for all used metagenomes are provided in Table S8. Metagenome assemblies generated for this study are available here: https://www.deemteam.fr/en/datasets. The corresponding proteomes as well as the HMM profiles are available in FigShare (https://figshare.com/projects/mVMRs/159701) and all code is available in GitLab (https://gitlab.com/anagtz/mvmrs).

## Author contributions

P.L.-G. and D.M. conceived the study and designed experiments, organized and participated in sampling and successive filtration steps, did the tangential filtration and quantitatively purified DNA from the different cell and viral fractions. A.G.-P. participated in sampling and carried out all bioinformatic analyses, with informatics support by P.D. M.K. designed the HMM profiles for MCPs. M.C. carried out flow cytometry counts of viruses and cells with the assistance of L.J. M.L.-P. and F.R.-V. co-organized sampling in marine and solar saltern sites and helped with filtration on board. P.L.-G. wrote the manuscript with input from A.G.-P. and M.C. All authors read, commented and approved the manuscript.

