## Supplementary Figures for "Metagenome-derived virus-microbe ratios across ecosystems"

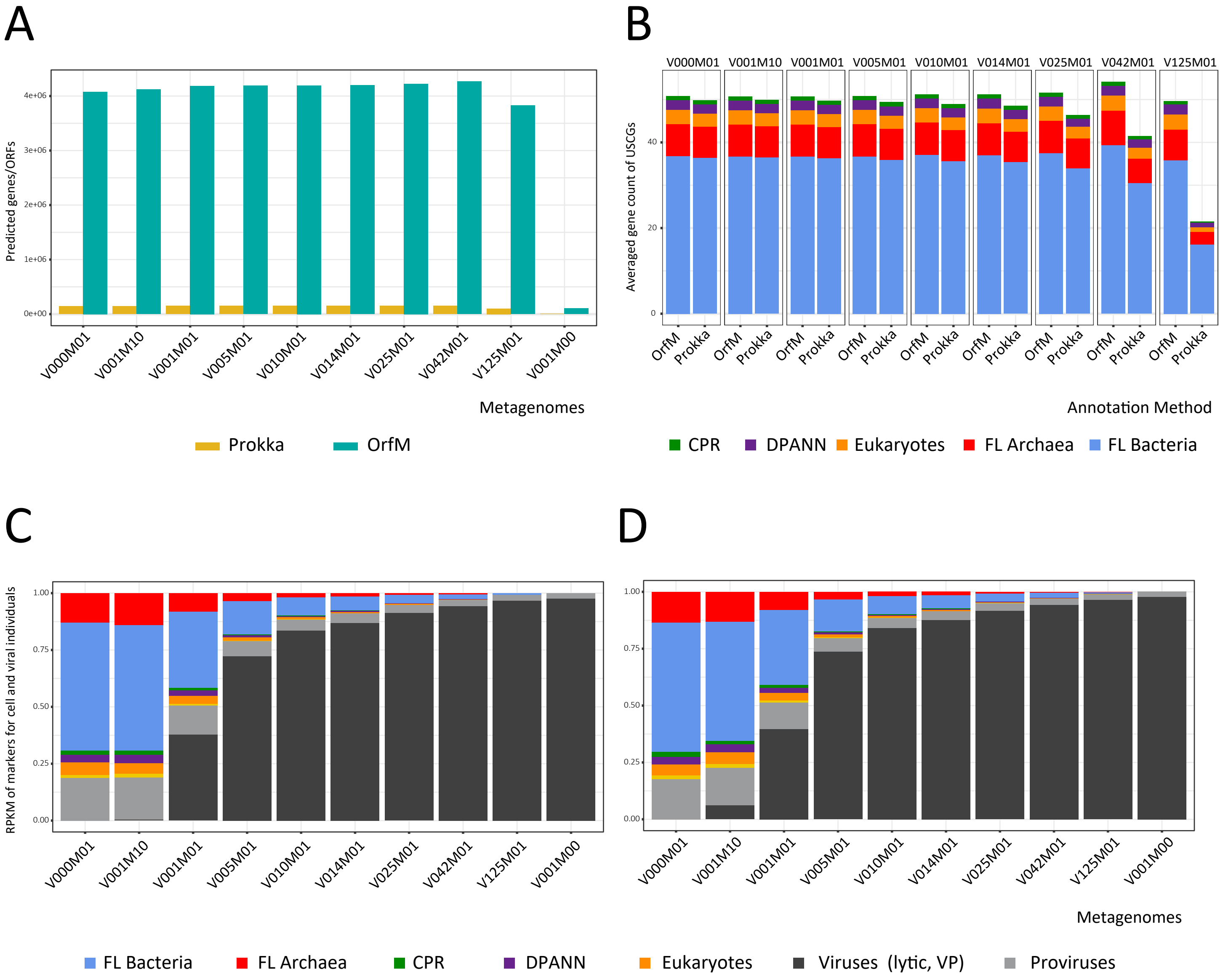


**Fig. S1. Predicted gene content of the generated mock metagenomes. A**, prediction of genes and open reading frames (ORFs) based on, respectively Prokka and OrfM annotation of assembled contigs. **B**, Prediction of the relative proportion of different cellular taxa in the mock metagenomes based on the averaged number of selected universal single copy genes (USCGs). CPR, Candidate Phyla Radiation, host-dependent epibiotic bacteria; DPANN, clade of host-dependent epibiotic archaea. **C**, Relative proportion of cellular and viral entities estimated from contig-based Prokka-annotated mock metagenomes with USCGs and major capsid proteins (MCPs) as markers for individuals. **D**, the same estimation for read-based OrfM annotation.

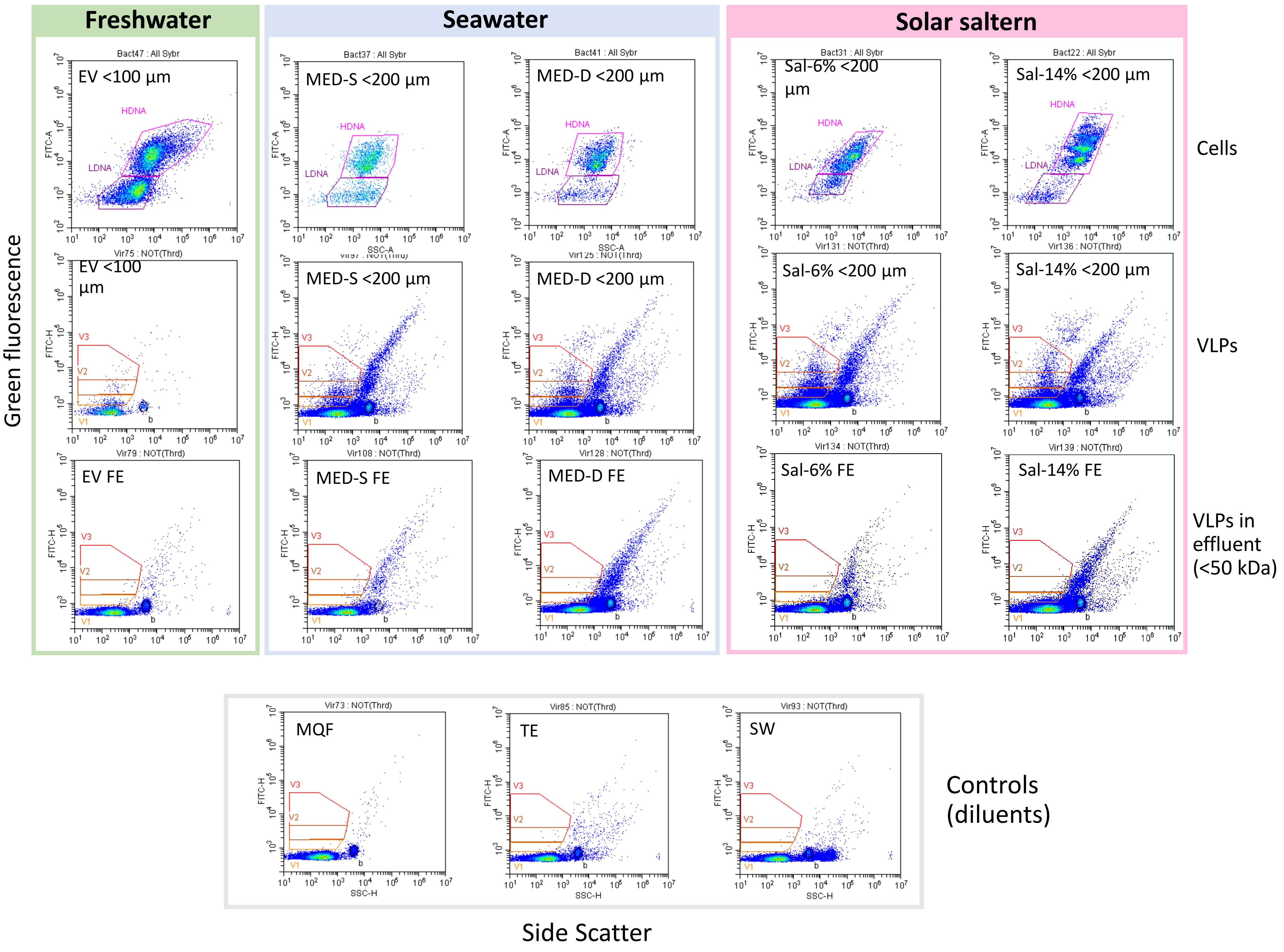


**Fig. S2. Cytograms of SYBR Green I-stained cells and virus-like particles (VLPs) in several aquatic environments spanning different salinities.** Data are shown for plankton below 200 µm cell size (100 µm for EV). Three VLP subpopulations (V1-V2-V3) and two major cellular subpopulations containing high DNA (HDNA) and low DNA (LDNA) levels are defined. Blanks are also shown: MQF, Milli-Q water, TE, Tris EDTA buffer, SW, seawater. FE, final eluate (<50 kDa); b, fluorescent 0.2µm beads.

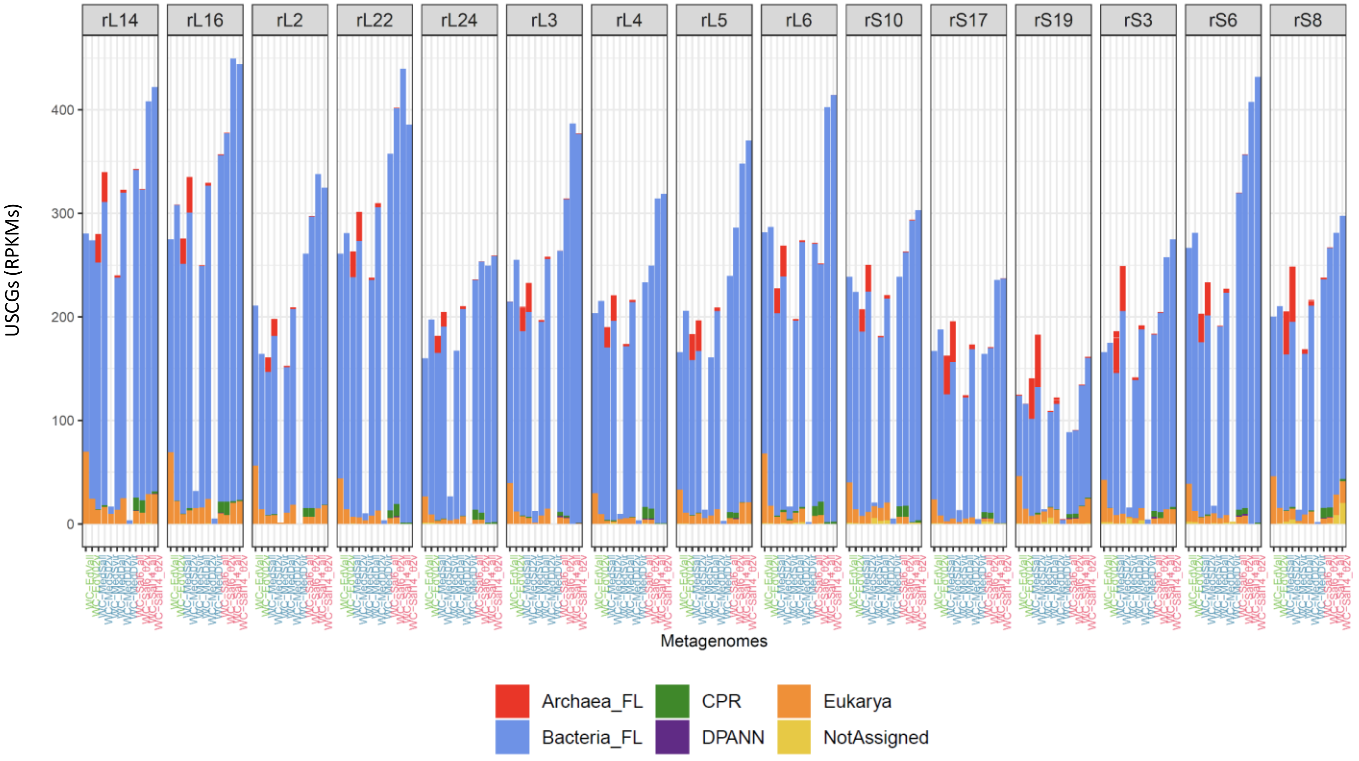


**Fig. S3. Normalised counts of selected USCGs in aquatic metagenomes of coexisting viral and cell fractions.** Suffixes in sample names indicate the plankton size fraction: _all, >50 kDa - 200 µm (100 µm in EdV); _02V, >50 kDa - 5 µm; _vir, >50 kDa - 0.2 µm ('metavirome'). Samples correspond, from left to right to plankton of the Etang des Vallées, France (EdV), Mediterranean water column, 20 m depth (MedS) and 40 m (MedD), and Bras del Port solar saltern ponds at 6-14% salt (Sal06, Sal14). Counts are expressed in reads per kb per million mapped reads.

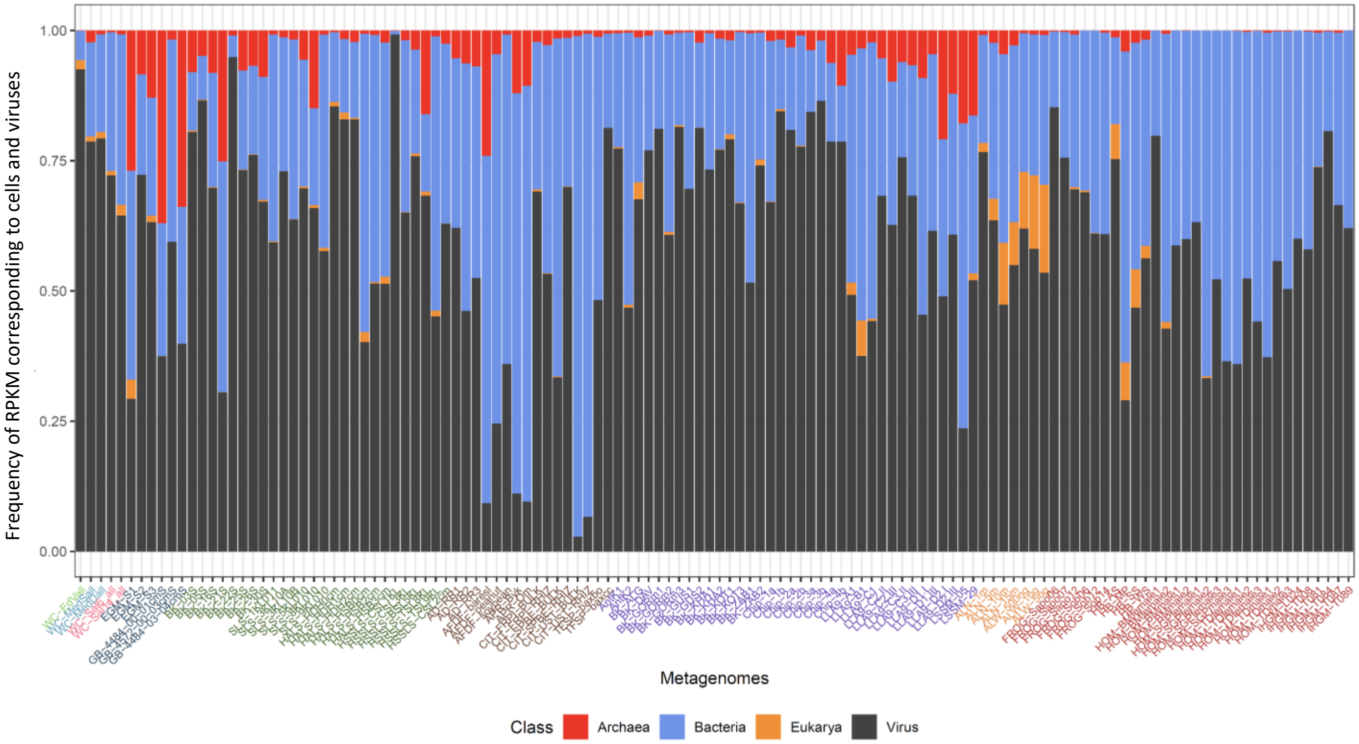


**Fig. S4. Relative abundance of cells and viruses across the studied metagenomes.** Cells were counted using the most represented USCG per metagenome.

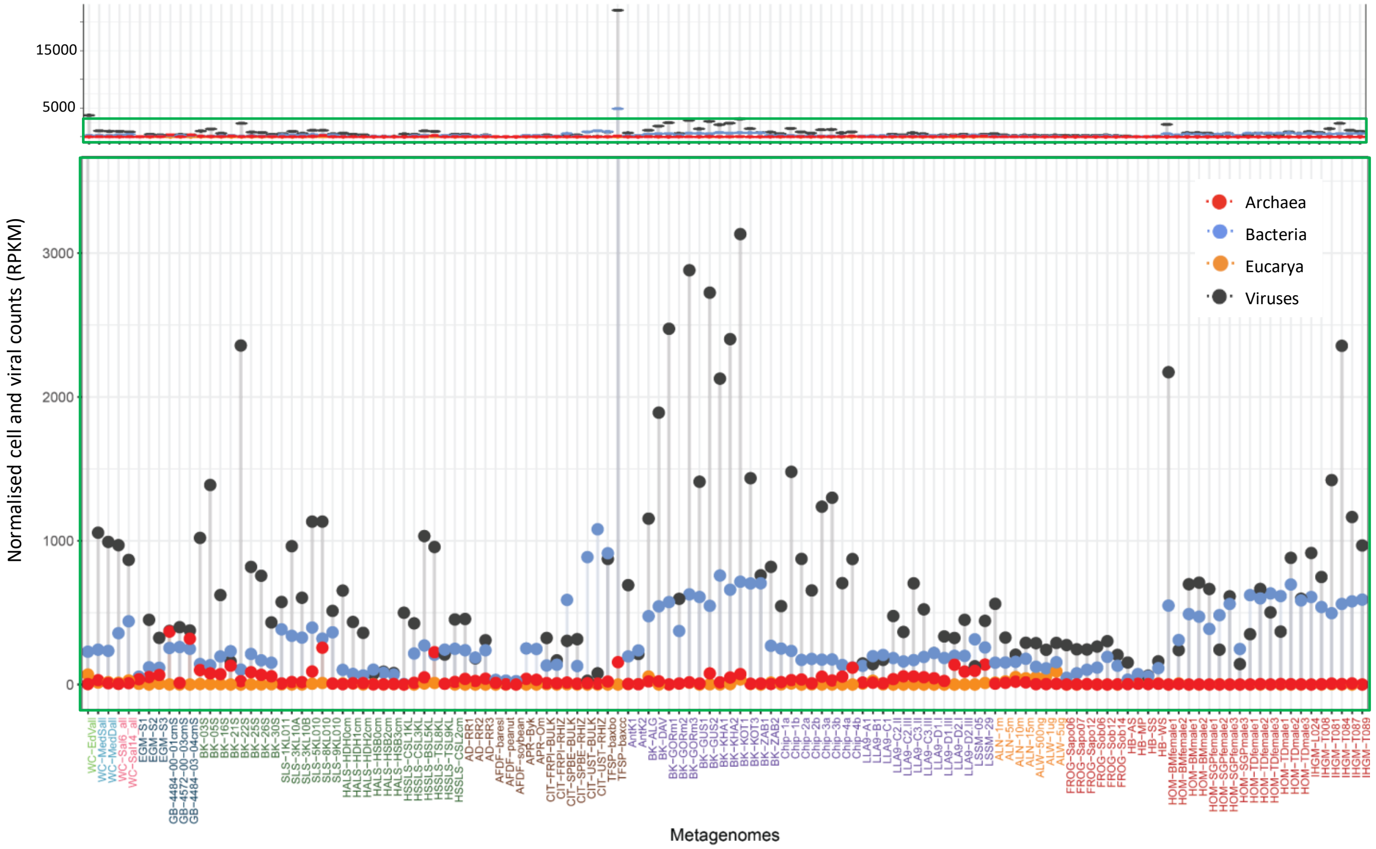


**Fig. S5. Normalised metagenome counts of viruses and microbial cells of the three domains of life across ecosystem types.** Viruses were counted using MCPs (sum of all) and cells for each large cellular group using the most abundant USCG for that group. All the metagenomes were analysed using the same pipeline from raw reads. RPKM, reads per kilo base per million mapped metagenome reads. The lower panel shows a zoom on the area covered by the green rectangle in the upper panel.

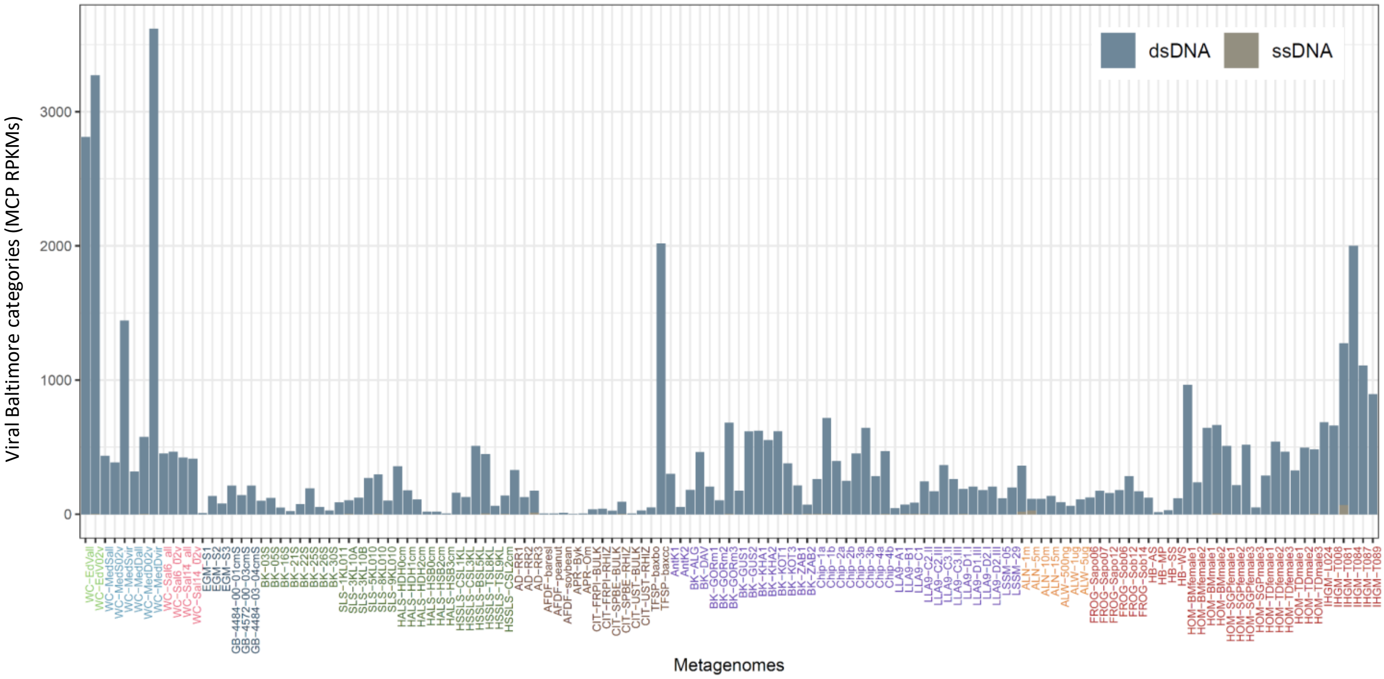


**Fig. S6. Relative proportions of viral Baltimore classes identified across metagenomes using major capsid proteins as markers.**

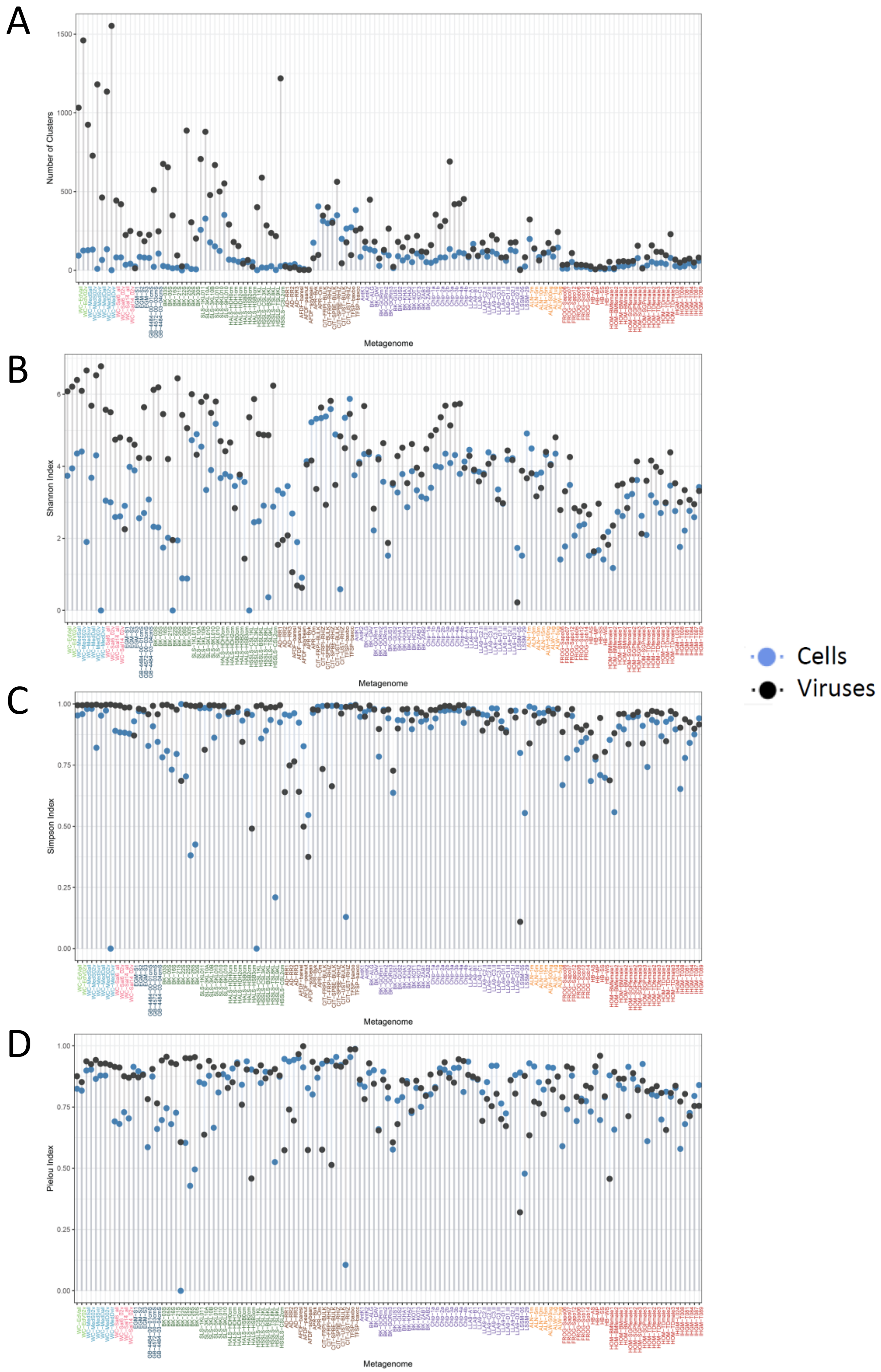


**Fig. S7. Diversity indices of viruses and cells across ecosystems. A**, Alpha diversity, corresponding the number of viral and cellular "species" clusters. **B**, Shannon index (diversity). **C**, Simpson index (diversity). **D**, Pielou index (evenness).

**A**

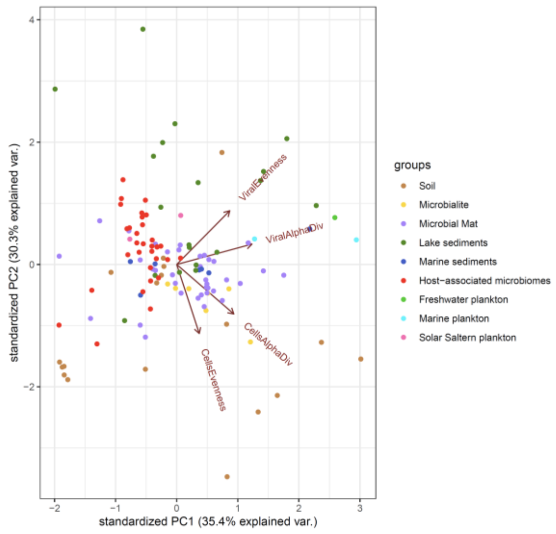


**B**

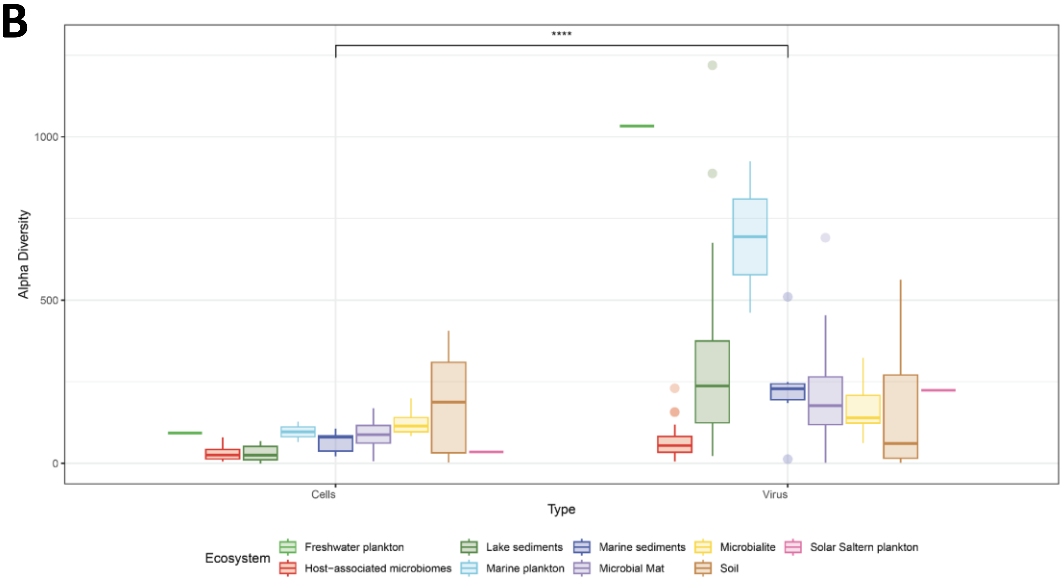


**Fig. S8. Diversity parameters of cells and viruses per ecosystem type. A**, Principal component analysis of different metagenomes maximizing the variance explained by viruses (MCPs counts in RPKMs) and microbial cells (USCG counts in RPKMs). Metagenomes are coloured as a function of ecosystem type. **B**, Boxplot showing the alpha diversity of cells and viruses per ecosystem type. Diversity differences between the cells and viruses are significant ($p<0.0001$).
